# Systematics of the oil bee genus *Lanthanomelissa* and its implications to the biogeography of southern grasslands

**DOI:** 10.1101/2020.03.12.989681

**Authors:** Taís M. A. Ribeiro, Aline C. Martins, Daniel P. Silva, Antonio J. C. Aguiar

## Abstract

*Lanthanomelissa* is a controversial taxonomic group of bees that was treated as subgenus of *Chalepogenus* or an independent genus. All of its species are endemic to the south-eastern grasslands of South America, an endangered and still poorly known environment. We aimed to understand the origin of this group of bees in time and space as well as the influence of Quaternary climatic fluctuations on its current distribution and possible link to the history of the Southern Grasslands. We inferred phylogenetic relationships of *Lanthanomelissa* species using 37 terminals and 3430 nucleotides of three mitochondrial and two nuclear markers and estimated divergence times and ancestral geographic range. We performed an ensemble with the algorithms SVM, Maxent, and Random Forest in a dataset of 192 georeferenced occurrence points using 19 WorldClim bioclimatic variables to analyse species distribution during the current and two past climatic scenarios (LIG, ∼120 kya and LGM, ∼21 kya). The results support the monophyly of the genus and taxonomic changes including the species *Lanthanomelissa parva* n. comb. on *Lanthanomelissa*, and the treatment of the *goeldianus* group of *Chalepogenus* as the genus *Lanthanella.* The genus originated at the Oligocene-Miocene border in the Chacoan-Pampean region, and the glacial-interglacial models indicated expansion in Last Glacial Maximum and retraction in Last Interglacial. Expansion and retraction of *Lanthanomelissa* distribution in the last glacial-interglacial indicated grasslands distributional shifts during periods of climate cooling and warming. The diversification of *Lanthanomelissa* supported the estimated expansion of southern grasslands in South America concurrently with the origin of Cerrado during the late Miocene. Their origin was approximately synchronized with their exclusive floral host, *Sisyrinchium* (Iridaceae).

## 1. Introduction

Bees are the most important angiosperm pollinators, relying entirely on flower resources (Vogel 1974; Michener 2007; Ollerton *et al.* 2011), with differing degrees of specialization (Fenster *et al.* 2004). Specialized bee-plant relationships tend to restrict their distribution according to their mutualist partner (Vamosi and Vamosi 2010). *Lanthanomelissa* bees are endemic to the Campos of southern Brazil, Argentina, and Paraguay, including the Pampa and Subtropical Highland Grasslands. The tribe where it belongs – the Tapinotaspidini – originated in early Palaeocene savannas and diversified in open vegetation environments until conquering forests in Eocene (Aguiar *et al.* 2020). Tapinotaspidini bees nest in the soil, as many other groups of bees who are intimately associated to open arid regions (Michener 1979), but they overcome the risks offered by humidity by the use of floral oils in nest lining (Aguiar *et al.* 2004).

The genus *Lanthanomelissa* is a controversial taxon of bees that has been treated either as a distinct genus with two subgenera, as a subgenus of *Chalepogenus*, or as an unique genus (Michener and Moure 1957; Michener 2007; Aguiar 2012). The classification of the two genera, *Chalepogenus* and *Lanthanomelissa*, in generic or subgeneric level was considered artificial by Michener (1963), and must be subject to further studies using phylogenetic methods.

The species of the oil-collecting bee genus *Lanthanomelissa* are solitary bees that nest on the ground close to populations of *Sisyrinchium* (Iridaceae) (Rozen *et al.* 2006), flowers in which they are specialists and resource-dependent (Cocucci and Vogel 2001). *Sisyrinchium* offer oil and pollen to these bees and about 35% of its species produce oil in trichomatic glands, while the remaining are pollen offering flowers and do not offer oil (Chauveau *et al.* 2011). The species in this genus are widely distributed in Neotropical Region, always in open vegetation environments, but its highest diversity is found around Parana Basin in Southern South America, overlapping with the distribution of *Lanthanomelissa* bees, their main pollinators (Cocucci and Vogel 2001; Chauveau *et al.* 2011; Chauveau *et al.* 2012). The closest relatives of *Lanthanomelissa* are part of the genus *Chalepogenus* (Aguiar *et al.* 2020) that explore plant species from the *Nierembergia* (Solanaceae) genus for oil and pollen (Cocucci 1991). The origin of *Sisyrinchium* and *Nierembergia* stem groups are close and estimated at about 13-20 Ma (Martins *et al.* 2015).

*Lanthanomelissa* and most of the oil-producing *Sisyrinchium* are endemic to Southern South American Grasslands. These grasslands comprehend a large open vegetation area occupying parts of southern Brazil, Argentina, and Uruguay included in two biomes: Atlantic Forest (Subtropical Highland Grasslands) and Pampa (Cabrera & Willink, 1973; IBGE, 2004; Overbeck et al., 2007; Morrone, 2014). Subtropical Highland Grasslands in Atlantic Forest occurs in patches forming a mosaic with *Araucaria* Forest at elevations from 700 to 1300 m a.sl. (Overbeck *et al.* 2015). The Pampa is part of the Chacoan biogeographic subregion, together with the biomes included in the South American Diagonal of Open Formations: Chaco, Cerrado and Caatinga (Morrone 2000). The origin and relations of the Pampa and Subtropical Highland Grasslands are still barely known. The Miocene climatic cooling, Andean Uplift, and the evolution of C4 grasses have been argued as the main factors influencing open vegetation worldwide and the formation of the Dry Diagonal in South America (Simon *et al.* 2009; Pennington and Hughes 2014).

The present biome configuration of forested and open vegetation remained relatively stable since the Pliocene (5.3 - 2.5 Ma). During the last 2.5 Ma in the Quaternary, however, fluctuations between global climate cooling and warming events (Hewitt 2004) could have led to periods of expansion and retraction of grasslands in south-eastern South America vegetation (Behling 1997; Safford 1999; Behling 2002; Behling *et al.* 2004; Safford 2007; Behling and Pillar 2007). Biogeographic studies with solitary bees (Zanella 2002; Ramos and Melo 2010), spiders (Ferretti *et al.* 2014), and birds (Porzecanski and Cracraft 2005) suggest that the Pampa is related to Chaco, Atlantic Forest, and Monte vegetation. This is reinforced by the occurrence of xerophytic elements from Chaco, like the cactus *Opuntia*, and patches of Atlantic forest (Boldrini 2009). Since bees are intrinsically connected to the vegetation, their origin and diversification could help understand biome evolution. In this context, we investigated the early origin and geographic distribution through time of the endemic bee genus *Lanthanomelissa*, through molecular phylogenetics associated to biogeography and distribution models.

We were interested in understanding the origins, distributions and persistence of these bees in south-eastern South America grasslands. Moreover, we were interested in finding out if *Lanthanomelissa* originate synchronously with its unique oil host *Sisyrinchium.* To answer those questions, we reconstructed the diversification of *Lanthanomelissa* in time using molecular phylogenetic tools coupled with ancestral range reconstructions and species distribution modelling, to find out when and under which conditions this bee group arose and how was the influence of recent climatic fluctuations on its distribution. We hypothesized that *Lanthanomelissa*’s origin, diversification and Quaternary fluctuations are directly associated to the origin of South America grasslands and that of its oil hosts, the plants of the genus *Sisyrinchium*.

## 2. Material and methods

### 2.1 Taxon sampling, DNA sequencing, and phylogenetic analysis

We analysed all five species recognized in *Lanthanomelissa* (sensu Urban 1995), with more than one sample per species in order to maximize the variation within the species and investigate each species dispersal history. We tried to sample all the known distribution of *Lanthanomelissa*, including high and lowland grasslands. The genera *Chalepogenus, Arhysoceble*, and *Trigonopedia* were used as outgroups, since they are the closest relatives to *Lanthanomelissa*, according to the comprehensive analysis for the Tapinotaspidini tribe by (Aguiar *et al.* 2020). We used specimens preserved in 95% EtOH, but also included pinned specimens collected less than 10 years ago for DNA extraction (Table S2). We tested several commercial DNA extraction kits (Table S2) following their recommended protocols. We adopted the following modifications for old museum specimens: Incubation in 70% EtOH for 24h for decontamination and rehydration; and incubation in Elution Buffer for 24h prior to digestion (Evangelista *et al.* 2017). We used different body parts aiming to improve the quality and quantity of DNA extracted (Table S2), maintaining the body integrity to assure further morphological studies.

We have newly generated 93 DNA sequences, plus 16 sequences from (Aguiar *et al.* 2020), all deposited in GenBank (Table S1). Vouchers are deposited in University of Brasilia’s Entomological Collection or in the collections listed in Table S1. We sequenced five genes: the mitochondrial genes cytochrome oxidase subunit I (COI, ∼700 bp), cytochrome B (CytB, ∼600 bp), and 16S (∼500 bp); and the nuclear long wavelength rhodopsin (LW-rhodopsin, ∼800 bp), and elongation factor 1-α F2 copy (EF-1α, ∼1100 bp). We chose both fast evolving mitochondrial genes and slow evolving protein coding genes to provide useful information as well as differentiation between species (Danforth *et al.* 2005; Danforth *et al.* 2013). Table S3 shows primers and PCR conditions. We conducted all extraction and amplification procedures in the Molecular Biology Laboratory at the Zoology Department of University of Brasilia and sent samples for purification and sequencing in the company Macrogen (South Korea). We assembled, trimmed, edited, and BLAST-searched the yielded sequences using the GenBank database on Geneious 8.1.9 (Kearse *et al.* 2012). We aligned sequences in MAFFT extension (Katoh and Standley 2013) on Geneious 8.1.9, using the following parameters: 200PAM/k=2 for the nucleotide scoring matrix; 1.53 for gap opening penalty and 0.123 offset value. For CO1 and CytB we used the algorithm G-INS-i, recommended for sequences with global homology. For the protein-coding nuclear Elongation-Factor 1-α and LW-Rhodopsin we used E-INS-i, recommended for sequences with multiple conserved domains and long gaps; and for 16S we used Q-INS-i, recommended for sequences with secondary structure. We made minor adjustments by eye.

We divided the data matrix by gene, codon and, for the nuclear markers, also by intron and exon and searched for best models and partition scheme in PartitionFinder (Lanfear *et al.* 2012), selecting with corrected Akaike Information Criterion. All schemes and models are available in Table S4. We performed Maximum likelihood tree searches in RAxML 7.4.2 (Stamatakis 2006) using the graphical user interface raxmlGUI 1.3 (Silvestro and Michalak 2012) and 1,000 non-parametric bootstrap replicates (Felsenstein 1985). We conducted Bayesian tree searches in MrBayes (Ronquist *et al.* 2012) using a matrix composed by 37 taxa and 3430 nucleotides with 10 million generations and four chains sampled every 1000 generations. We used 14 dataset partitions, as yielded by PartitionFinder 2.1.1 (Lanfear *et al.* 2017) searches, with models selected by Corrected Akaike Information Criterion (AICc) using the greedy algorithm (Lanfear *et al.* 2012) (Methods S1, Table S4). We discarded the first 25% of trees as burnin. We analysed the Convergence of Markov Monte Carlo chains (MCMC) in Tracer (Rambaut *et al.* 2018). We edited all trees in FigTree 1.4.3 (Rambaut 2016).

### 2.2 Divergence times estimates

Since there is no known fossil for Tapinotaspidini we used a secondary calibration point from the widely sampled phylogeny of Tapinotaspidini tribe (Aguiar *et al.* 2020). The age of most recent common ancestor of *Lanthanomelissa* + *Arhysoceble* + *Chalepogenus* is 28 Ma (95% HPD 22.98-34.89). We therefore applied a normal distribution prior (with values of mean: 28, stdev: 4.0).

We estimated the divergence times for *Lanthanomelissa* in BEAST version 1.8.4 (Drummond *et al.* 2012) via the CIPRES server (Miller *et al.* 2010), relying on the same 37 taxa matrix. We estimated the divergence times for *Lanthanomelissa* with the uncorrelated lognormal relaxed model. We used the Yule tree speciation model, which is more appropriate when considering sequences from different species (Heled and Drummond 2012; Drummond *et al.* 2015) and HKY substitution model with empirical base frequencies. We ran a Markov chain Monte Carlo (MCMC) for 50 million generations sampled every 10000 generations. We assessed convergence of chains in Tracer vs. 1.6 (Rambaut *et al.* 2018), considering an Effective Sample Size (ESS) for all parameters > 200. We produced the maximum clade credibility tree in TreeAnotator vs. 1.8.4 (part of BEAST package), with a burnin of the first 25% trees. We visualized and edited the trees in FigTree vs. 1.4.3 (Rambaut 2016).

### 2.3 Ancestral Area Reconstruction

We derived six biogeographic areas from the classification of Neotropical Region defined in Morrone (2014), representing the geographic range of *Lanthanomelissa* and outgroups: A) Atlantic, B) Cerrado, C) Chaco, D) Caatinga, E) Pampa, F) *Araucaria* Forest. For outgroups, we coded the geographic areas representative for all species of the lineages. For ancestral area estimation we relied on the R package ‘BioGeoBEARS’ (Matzke 2013), which evaluates the contribution of evolutionary processes (i.e., range expansion, range extinctions, vicariance, founder-event speciation) through the biogeographic models: DEC (Ree and Smith 2008), and modified versions of DIVA (Ronquist 1997), and BayArea (Landis *et al.* 2013) named DIVA-like and BayArea-like. We do not implement DEC+J model since it has been demonstrated to be a poor model of geographic range evolution (Ree and Sanmartín 2018). We conducted likelihood ratio tests based on AICc scores to assess the fitness of the models.

We coded the geographic area according to the vegetation type of the collection locality, even when representing a minor patch of vegetation located inside a biome. For instance, although the classification from Morrone (2014) suggested the Atlantic province covering all the costal Santa Catarina and highlands of São Paulo states, we considered the records on Cotia (São Paulo) and Maracajá (Santa Catarina) as occurring in patches of grasslands in the Atlantic Forest, and have coded them as *Araucaria* Forest Province. We did the same for records in Parana province. For discussion purposes we also consider the regionalization introduced by Olson et al. (2001) as modified by Antonelli et al. (2018).

### 2.4 Occurrence and environmental data sampling

We obtained 192 georeferenced points (Table S6) for all *Lanthanomelissa* species from literature and voucher labels deposited in entomological collections (Methods S1). When geographic coordinates were not available on the labels, we used Google Maps (Google Inc. 2018) to acquire approximate geographical coordinates from city downtown. We have also obtained data from the online databases SpeciesLink (www.splink.org.br) and GBIF (Global Biodiversity Information Facility, www.gbif.org/). The following entomological collections provided geographic location points for *Lanthanomelissa*: Departamento de Zoologia da Universidade de Brasília (DZUB), Departamento de Zoologia da Universidade do Paraná (DZUP), Museu de Ciências e Tecnologia da PUCRS (MCTP), Fundação Zoobotânica do Rio Grande do Sul (FZB/RS), American Museum of Natural History (AMNH), Coleção Sersic-Cocucci, Departamento de Botânica da Universidade Nacional de Cordoba (UNC), Museu Argentino de La Plata – Universidade Nacional de La Plata (UNLP), Universidade do Extremo Sul Catarinense (UNESC), Faculdade de Ciências e Letras de Ribeirão Preto USP - Coleção Camargo (RPSP), Museu de Zoologia da Universidade de São Paulo (MZSP). We have compiled 170 unique points, being 52 for *L. betinae*, 48 for *L. clementis*, 35 for *L. discrepans*, 21 for *L. magaliae*, and 14 for *L. pampicola*.

To estimate potential geographic range of *Lanthanomelissa* species across a glacial-interglacial cycle, we performed species distribution models based on current (from the years 1970 to 2000), Last Glacial Maximum (LGM, 22 ky ago) and Last Interglacial (LIG, 120 ky ago) bioclimatic variables from WorldClim (Hijmans *et al.* 2005; Otto-Bliesner *et al.* 2006; Fick and Hijmans 2017) in 2.5 arcminutes resolution.

For the Last Glacial Maximum and current climatic scenarios, we performed the simulations based on the Community Climate System Models (CCSM) general circulation model in 2.5 arcminutes resolution. To correspond to the other environmental layers, we resampled map resolution for Last Interglacial variables from 30 arcseconds to 2.5 arcminutes using the ‘raster’ package (Hijmans 2017) in R 3.4.2 (R Core Team 2017), implemented in RStudio 1.0.153 (RStudio Team 2016). To avoid collinearity and model overfitting (Jiménez-Valverde *et al.* 2011) we standardized the 19 bioclimatic variables by subtracting the mean value for each cell and then divided this result by the standard deviation, so that all variables vary from −1 to +1 and have average equal to zero and variance equal to one. We then ran a Principal Component Analysis (PCA) creating new orthogonal spatialized principal components (PCs) to remove variable collinearity. We performed this analysis first for current climate and then projected its linear coefficients into the past climates (LGM and LIG) so that the PCs generated for the past were dependent to the current scenario.

Since the models consider only climate data, they do not acknowledge biotic factors or species dispersal ability (Soberón 2007), so we restricted the modelled area using maximum and minimum latitudes and longitudes from the occurrence points as parameters for the extent area modelled (longitude: minimum −70 and maximum −45; latitude: minimum −40 and maximum −20).

### 2.5 Species distribution modelling

We performed all ecologic niche modelling analysis with the R package ENMTML (Andrade *et al.* 2020). Methods S1 provide information about occurrence data and environmental data sampling. We partitioned *Lanthanomelissa* species occurrence dataset with checkerboard partition, in which data was divided in two subsets (50% each). The first subset produced the potential distribution and the second evaluated it and vice-versa. Then, we performed an ensemble through a consensus from algorithms that had TSS values above mean with three algorithms: the machine learning algorithms Maxent (Maximum Entropy, Phillips et al., 2004), Random Forest (Breiman 2001) and SVM (Support Vector Machines, Schölkopf et al., 2001; Tax & Duin, 2004). We evaluated model performance based on the AUC (Area Under the Curve, Allouche et al., 2006) values, which is a threshold independent matrix varying from 0 to 1 and considered good when above 0.8. We have also evaluated True Skill Statistics values (TSS, Allouche et al., 2006), a threshold-dependent metric varying between −1 to +1 in which values above 0.5 are acceptable and values above 0.7 are considered good. To estimate stable areas for each species, we multiplied the binary rasters yielded by the ensemble with all models from the three scenarios (LIG, LGM, and current) to find the intersection between them and plot them together.

## 3 RESULTS

### 3.1 Molecular Phylogeny

The aligned data matrix comprised 109 sequences and 3,430 nucleotides, in which 88 sequences belong to *Lanthanomelissa* specimens and 21 to the outgroup. Bayesian and Maximum Likelihood inference trees both recovered similar highly supported clades (Figure S2, Figure S3) The trees show *Arhysoceble* as sister to a clade containing *Chalepogenus goeldianus, Chalepogenus parvus* and *Lanthanomelissa* (Bayesian Posterior Probability (BPP) = 1 and bootstrap support (BS) = 97). *Lanthanomelissa* species are also all well supported. In Bayesian tree, *C. goeldianus* is sister to *C. parvus* + *Lanthanomelissa* (1 BPP) and *C. parvus* is sister to *Lanthanomelissa* (0.53 BPP). However, in Maximum Likelihood tree these relations are inverted as *C. parvus* is sister to *C. goeldianus* and *Lanthanomelissa* (97% BS) while *C. goeldianus* is sister to *Lanthanomelissa* (55% BS). *Lanthanomelissa* constitute a monophyletic group (81% BS, 0.99 BPP) and all the species are well supported as monophyletic (BS > 88%, BPP > 0.95). In this clade, *L. betinae* is sister to all the other species (81% BS, 0.99 BPP). In both trees *L. discrepans* and *L. magaliae* are monophyletic (73% BS, 0.97 BPP). These two species constitute a clade sister to another containing *L. pampicola* and *L. clementis* (87% BS, 1 BPP), however in Maximum Likelihood tree these two species are weakly supported as sister groups (54% BS) while in Bayesian tree this relation is highly supported (0.97 BPP).

### 3.2 Divergence times and ancestral range reconstruction

According to the Bayesian time tree (Figure 1, Figure S3), the crown group *Arhysoceble* + *Chalepogenus* + *Lanthanomelissa* had its origin estimated in the Oligocene, at 26.91 (18.56 – 34.68, 95% HPD) Mya. The crown group of *Chalepogenus parvus*+ *Lanthanomelissa* had its age estimated at 20.91 (13.24– 28.38) Mya, in Miocene. Crown age for *C. parvus* + *Lanthanomelissa* is estimated at 17.47 (11.16– 24.79) Mya. Crown age for *Lanthanomelissa* is estimated at 14.31 (8.58– 20.56) Mya. Individual *Lanthanomelissa* species had their crown ages estimated between 6.14 (2.59 – 10.16, 95% HPD for *L. pampicola*) and 3.12 (1.07 – 6.01 for *L. magaliae*).

**Figure 1.**
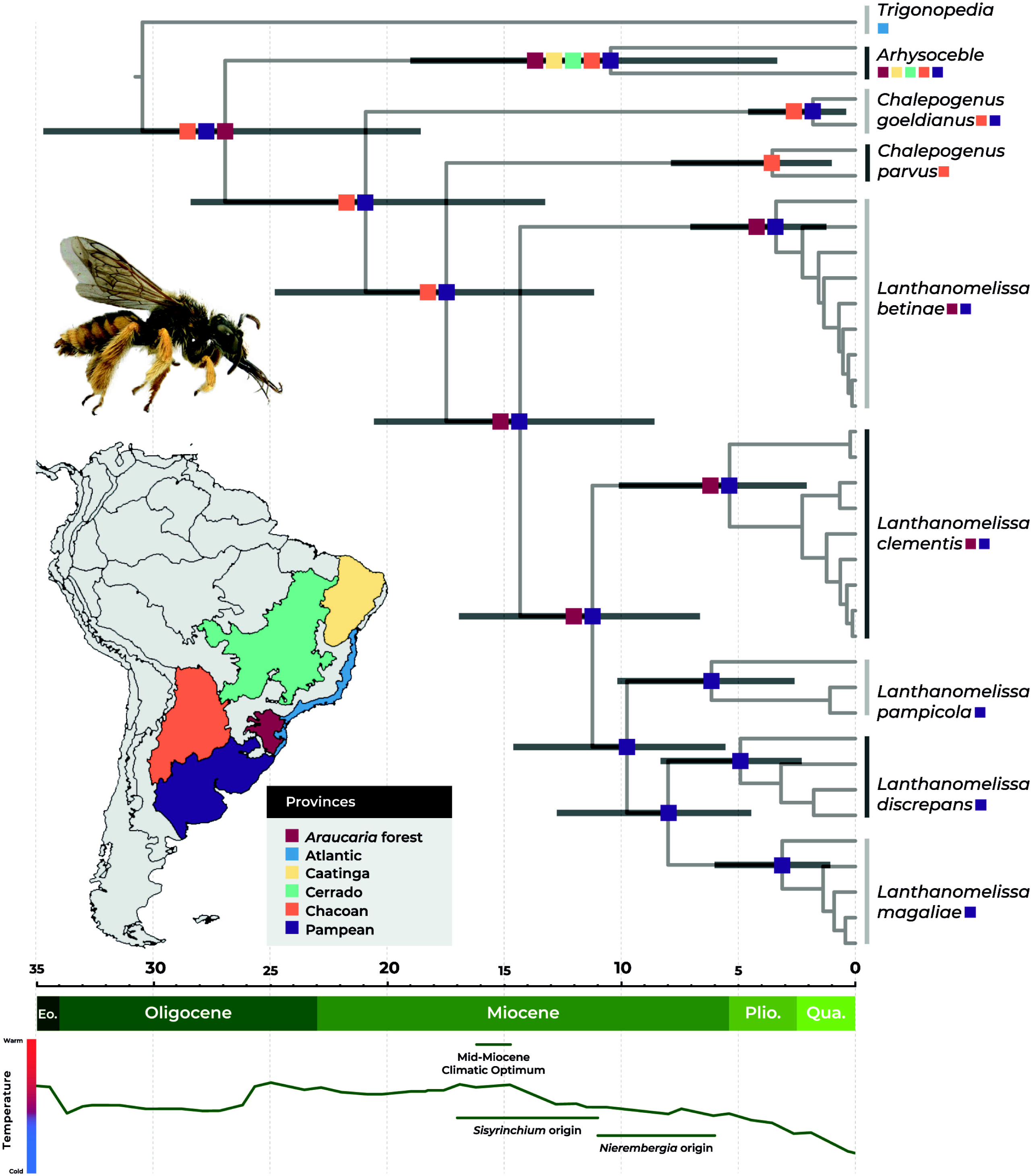
Time calibrated phylogeny and historical biogeography of *Lanthanomelissa* with representatives of *Arhysoceble, Chalepogenus* and *Trigonopedia* as outgroup based on a concatenated matrix comprising 37 terminals and 3430 nucleotides with 50 million generations, age of the root node sampled from a normal distribution (mean =28 and s.d. = 4). Map shows provinces from Morrone (2014) used for biogeographic reconstruction. Coloured squares after species names and at nodes represent current historical distributions inferred in BioGeoBEARS under the Bayarealike+j model (lnL = −28.65). Horizontal grey bars at nodes indicate 95% HPD of estimated divergence times and bottom bar indicates epochs and Quaternary period. Abbreviations: Eo: Eocene, Plio: Pliocene; Qua: Quaternary. Picture shows a pinned specimen of *Lanthanomelissa discrepans*. Bottom: Temperature variation for the last 35 My adapted from (Zachos et al., 2001), indicating Mid-Miocene Climatic Optimum and estimated origin of *Sisyrinchium* and *Nierembergia* (95% hpd, Martins et al., 2015).

Multi-model analysis in BioGeoBEARS yielded BayArealike including founder-event speciation event *j* as the best fitting model for our data, as it showed the higher log likelihood and lowest AICc (LnL = −29.95, AICc = 64.24, Table S5). According to this analysis (Figure 1), the ancestral area for the most recent common ancestor of the lineage *Arhysoceble + C. goeldianus + C. parvus* comprised Chacoan, Pampean and *Araucaria* Forest provinces. However, for the most recent common ancestor of the two *Chalepogenus* + *Lanthanomelissa*, the area comprised only Chacoan + Pampean provinces. For *Lanthanomelissa* the ancestral area inferred is *Araucaria* Forest + Pampean, and this is the current area for *L. betinae* and *L. clementis*, while the other three species are limited to Pampean province.

### 3.3 Distribution modelling of Lanthanomelissa

Figure 2 depicts the distribution modelling for all *Lanthanomelissa* species for current climate, Last Interglacial, Last Glacial Maximum, and stable areas. Figure S5 shows separately each modelling results and all statistic values are listed in Table S7. Potential distribution of *Lanthanomelissa* species based on current climate indicates that all species could occur in larger areas of grassland vegetation, but either collecting effort was not yet exhaustive or other factors, such as mutualistic partner, limits their distribution (see blue areas in Figure 2). For all *Lanthanomelissa* species the distribution modelled for the Last Interglacial (LIG) extended to the south, including mostly southern Brazil and Uruguay, almost reaching the latitude of −40 (Figure 2). On the Last Glacial Maximum (LGM) this suitability was shifted northwards. For all species in LGM, the suitability extrapolated current continental borders because the sea level was retracted at that time.

**Figure 2.**
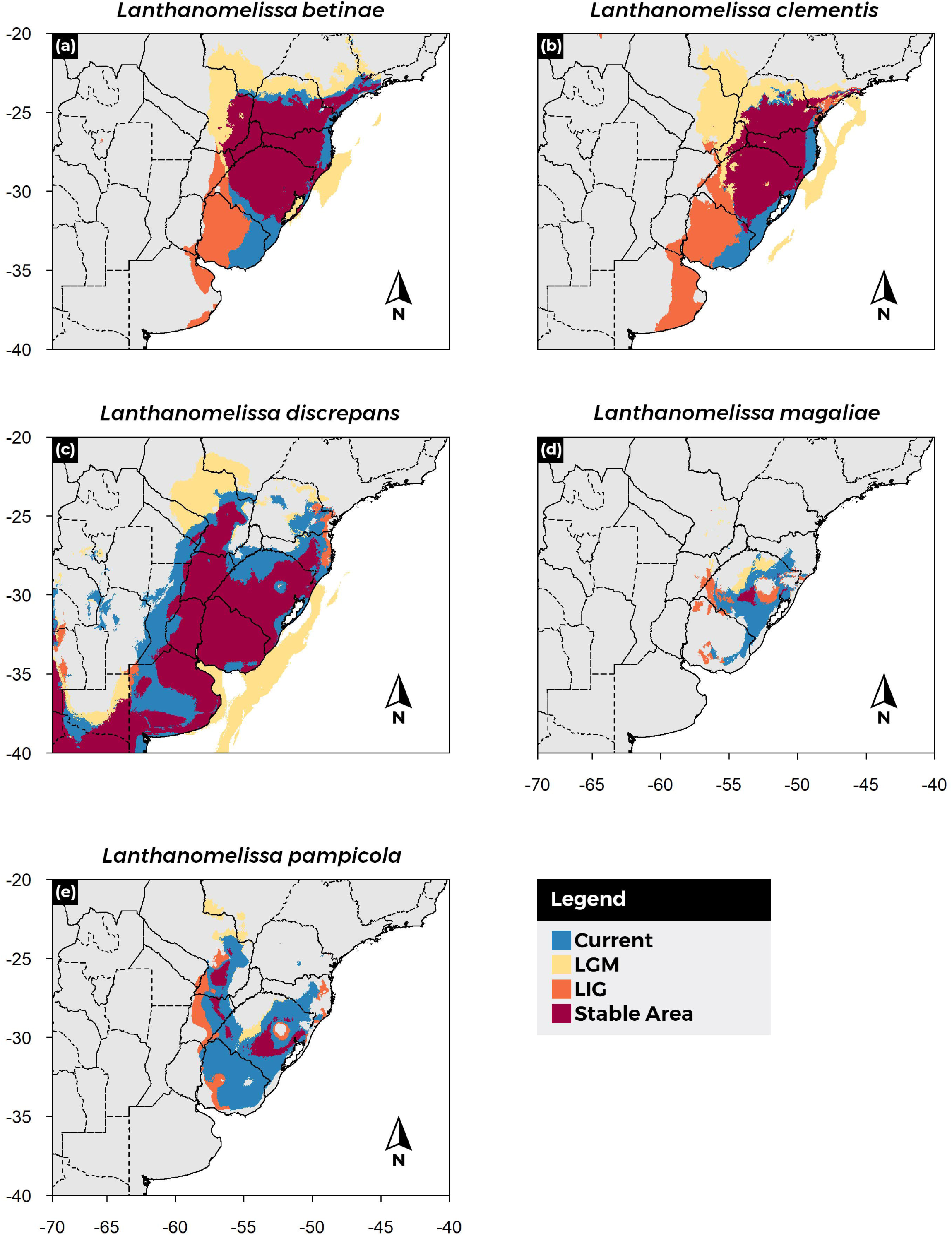
Potential range shifts for *Lanthanomelissa betinae* (a), *L. clementis* (b), L. *discrepans* (c), *L. magaliae* (d) and *L. pampicola* (e). Stable areas are shown in purple, areas predicted as suitable in the Last Interglacial (LIG) are shown in orange, areas predicted in the Last Glacial Maximum (LGM) are in yellow and areas in current distribution are shown in blue.

Stable areas predicted for *L. betinae* and for *L. clementis* (Figure 2 a, b) included all three South Brazilian states (Paraná, Santa Catarina, and Rio Grande do Sul) and even a small part of São Paulo. For these two species, however, the stable areas included only a small portion of the Pampa. For *L. discrepans* (Figure 2c) the stable area is larger than for the other species, including parts of eastern Argentina, and probably expanding southwards pass latitude −40S. Northwards, it reaches part of Paraguay and a small portion of the Brazilian state of Santa Catarina. Stable areas predicted for *L. magaliae* and for *L. pampicola* (Figure 2 d, e) are very restricted in relation to the other species, limited to only a small portion of Rio Grande do Sul for *L. magaliae*, while for *L. pampicola* it includes patches in Argentina and Paraguay.

## 4 DISCUSSION

### 4.1 Phylogenetic relationships and taxonomic considerations

Our phylogenetic results indicated *Lanthanomelissa* as a monophyletic genus sister to species of the paraphyletic genus *Chalepogenus* (Aguiar *et al.* 2020). The species currently described in *Chalepogenus* form two paraphyletic groups: the most speciose are composed mostly by species occurring in the Monte vegetation in Argentina (*Chalepogenus* s.s.); the remaining *Chalepogenus* are related to *Lanthanomelissa*, related to Pampas and Southern grasslands, as shown in the present paper. This result reinforces the need of a taxonomic reclassification of the species of *Chalepogenus* related to *Lanthanomelissa*, as we presented below. Additionally, the species grouped in the *Chalepogenus* s.s. should be subject of further taxonomic review.

As indicated in our phylogeny (Fig 1, S1 and S2) *Chalepogenus goeldianus* is sister to all *Lanthanomelissa* plus *Chalepogenus parvus*. Therefore, we propose that *C. goeldianus* and related species (not sampled in our phylogeny) must be reclassified in *Lanthanella* Michener & Moure (1957). *Lanthanella* was proposed as subgenus of *Lanthanomelissa* by Michener and Moure (1957) as monotypic for *L. (L.) goeldiana*, and posteriorly Moure (1992) surged the group to genus level. Urban (1995) described *Lanthanella luciane* Urban 1995, and Roig-Alsina (1999) transferred *L. goeldiana* and *L. luciane* to *Chalepogenus*, also describing a new species to this group, which he named *Chalepogenus neffi*. These three species were proposed by Roig-Alsina (1999) as goeldianus species group and the present results support the significance of this clade as the distinct genus *Lanthanella* therefore composed by *Lanthanella goeldiana* (Friese, 1899), *Lanthanella luciane* Urban, 1995, and *Lanthanella neffi* n. comb. Roig-Alsina (1999) and reinforced by morphological characters such as identical genital capsules, lower apices of hind tibia acute (Roig-Alsina 1999),

Due to their sister position to *Lanthanomelissa*, we propose *Lanthanomelissa parva* n. comb., instead of designating a new genus. As a result, *Lanthanomelissa* is now composed by six species, supported also by morphological characters such as the two submarginal cells in the fore wing, reduced body size, and similar terminalia of males.

### 4.2. Lanthanomelissa origin indicates the presence of grasslands in South America in early Miocene

The origin of the stem group *Lanthanomelissa* was estimated at the transition from Oligocene to Miocene (20.91, 13.24 – 28.38 Mya) in an ancestral area comprising Pampean and Chacoan biogeographical provinces. Further, these two areas could have been fragmented by the Patagonian Sea, an ocean transgression extended from southern Patagonia to Bolivia and facilitated by Andean orogeny process (Ortiz-Jaureguizar and Cladera 2006). The crown group *Lanthanomelissa* originated at 14.31 (8.58 – 20.56 Mya) in the Middle Miocene Climatic Optimum, which was the peak of a global warming phase that started in late Oligocene (Zachos *et al.* 2001) and favoured the raise of many animal and plant groups specially in South America (Hoorn *et al.* 2010).

The origin of the crown group *Lanthanomelissa* is associated to the beginning of a phase of climatic cooling and expansion of open vegetation environments in many parts of the world, including South America. From 15 My Andean Cordillera elevations created a rain-shadow effect to humid winds from west increasing the aridity in eastern South America (Hartley 2003). The higher aridity associated to the C4 grasses expansion led to the predominance of open vegetation habitats in southern South America (Lehmann *et al.* 2011; Strömberg 2011; Werneck 2011; Hoetzel *et al.* 2013; Pennington and Hughes 2014). Moreover, in late Miocene, the low CO_2_ pressure coupled led to a worldwide expansion of C4 grasslands, more flammable than common grasses, and the origin of savannas (Pennington and Hughes 2014), which rise to dominance in the Pliocene (Simon *et al.* 2009). Similarly, the origin of Pampa inferred here at ca. 10 Mya could have been a result from the expansion of the grasslands associated with *Araucaria* forest. This suggests that the Cerrado and the Pampa could have gone through similar diversification processes associated to the expansion of grasslands. The expansion of grasslands could have also influenced the origin of the clade *L. pampicola* + *L. discrepans* + *L. magaliae* already in a Pampa area, differently from *L. betinae* and *L. clementis*, which maintained their distribution in the *Araucaria* forest province.

### 4.3 Synchronous origin of oil source plants and Lanthanomelissa bees

*Lanthanomelissa* crown age is coincident with the stem origin of its preferred oil host, plants species from the *Sisyrinchium* genus, as well as is the origin of the stem node, thus including *Chalepogenus parvus and C. goeldianus*, coincident with *Nierembergia* (see Martins et al., 2015 for *Nierembergia* and *Sisyrinchium* ages). The coincident age, spatial distribution, and close mutualistic association (Cocucci and Vogel 2001; Chauveau *et al.* 2011) supports our suggestion that *Lanthanomelissa* and *Sisyrinchium* could have played important roles in each other’s diversification and biogeographic history in South America. The gains of glandular trichomes in species of *Sisyrinchium*, especially the oil secreting ones, have been hypothesized as one of their main drivers of diversification in South America (Chauveau *et al.* 2011). Although the presence of oil glandular trichomes is not ancestral in *Sisyrinchium*, it occurred in early diverging clades (Chauveau *et al.* 2011), suggesting an ancient relationship with oil bee pollinators. Further analysis on diversification of these plant clades is necessary to understand the role of oil bee pollinators in speciation or extinction. *Lanthanomelissa*’s sister lineages, *i.e.* lineages of *Chalepogenus* and *Arhysoceble*, would have exploited other plants for oils, as demonstrated by time of origin we estimated, current preference for oil hosts (Cocucci 1991; Renner and Schaefer 2010; Martins and Alves-dos-Santos 2013), and estimated origin of oil producing Solanaceae and Plantaginaceae (Martins *et al.* 2014; Martins *et al.* 2015). The relationship of the lineage *Arhysoceble*-*Lanthanomelissa* with floral oil families is intimately associated to the expansion of open vegetation areas in South America (Aguiar et al. in prep). The origin of floral oil glands in these younger plant clades evolved as an exaptation of the oil-bee-syndrome evolved with the Malpighiaceae since Palaeocene (Aguiar et al. in prep; Renner & Schaefer, 2010).

### 4.4 Climatic fluctuations drove Lanthanomelissa distribution towards grasslands areas in the Quaternary

According to our species distribution modelling, the suitability for most species shifted north- and eastwards in the LGM. This is corroborated by the presence of a mosaic of grasslands and *Araucaria* forest in the coast of southern Brazil, from São Paulo to Rio Grande do Sul (Behling 2002; Behling *et al.* 2004; Overbeck *et al.* 2007; Pessenda *et al.* 2009; Leite *et al.* 2016; Silva *et al.* 2018). The distribution of *L. discrepans* and *L. pampicola* are divided by a central gap suggesting the division of inland western and eastern clusters, as well as a costal cluster in LGM. This structure was also observed in this area for plants endemic to the Southern Grasslands (Pinheiro *et al.* 2011; Longo *et al.* 2014; Silva *et al.* 2018). The species could have dispersed through the “Portal de Torres” (Rambo 1950), a migratory route from the coastal Atlantic forest to the inland grasslands (Pinheiro *et al.* 2011; Barros *et al.* 2018; Silva *et al.* 2018).

At LGM, grasslands were dominant in south eastern South America and climate was drier and colder than in the present (Behling 1997), and possibly extended to the coast of Uruguay, as this area is also predicted as suitable for *Lanthanomelissa*. About 4 Kya, with increasing temperature the *Araucaria* Forest expanded, limiting once again the occurrence of the grasslands (Behling *et al.* 2004), as well as the distribution of *Lanthanomelissa*, as evident in models for current distribution of *L. clementis.* The fluctuation among grasslands and *Araucaria* forests have been suggested as important drivers of speciation in Subtropical Highland Grasslands, but not in Pampa where registers for *Araucaria* Forest are too recent (Fregonezi *et al.* 2013). Instead, abiotic factors (e.g. soil diversity) and ecological relations (e.g. intraspecific competition and pollination) could have been stronger selective pressures in this region. However, few studies use species that cohabit both environments to address the interaction between these abiotic factors and how the two grassland areas relate (Fregonezi *et al.* 2013; Peres *et al.* 2015).

The Last Interglacial (LIG, ∼120 kya) had the highest global temperatures of the last 250 ky and is characterized by wet conditions, higher sea levels and expansion of forest biomes over arid open formations (Otto-Bliesner *et al.* 2006). This could have caused the shown restriction on the distribution of most species from *Lanthanomelissa* at this time (Figure 2), suggesting a restriction in the grasslands, which were probably more expressive towards the south.

The stable areas indicated by *Lanthanomelissa* species in the last glacial-interglacial cycle varies among species, but a clear pattern of stability in highland grasslands can be observed in *L. betinae* and *L. clementis*. In colder periods, the highlands act as refugia to those species, since it favoured expansion of grasslands (Behling 1997). Other species respond differently to stable areas analysis, but we can see an overlap between stable areas that approximately correspond to the Southern Atlantic Forest refugia described in Costa et al. (2018). This region has been dynamic over climatic fluctuations of Quaternary, presenting forest refugia in the grassland phase, which rapidly expanded over grasslands when the climate became suitable (Costa *et al.* 2018).

The potential distribution based on current climate indicated large areas as suitable for most *Lanthanomelissa* species, including areas of grasslands and Chaco. Collecting effort has been not yet exhaustive, but possibly other factors, such as *Sisyrinchium* plants distribution limits their distribution.

*Lanthanomelissa* are dependent on their flowers, although the contrary is not always valid (Cocucci and Vogel 2001; Chauveau *et al.* 2012). However, the congruence of the diversification of those bees with the origin of its oil host indicates that those groups could have been drivers of diversification to each other. Integrative modelling approaches using occurrence data of the oil-host plant *Sisyrinchium* as a variable limiting the distribution of *Lanthanomelissa* and the phylogeography of one of these bee species would also bring valuable insights on the biogeographic history of the south-eastern South America grasslands. Even so, understanding the systematics and the evolutionary history of bees can help understand biome evolution, as the origin of the genus *Lanthanomelissa* in an ancestral area shared by Pampa and Chaco indicates that it is related to the origin of the South America Grasslands.

## Data availability statement

Specimen voucher information and geographic localities used in this study are provided as Supporting Information (Table S1 and S6 respectively). Genetic sequences were deposited in GenBank and accession numbers are provided in Supporting Information (Table S1). Sequence matrix, phylogenetics trees and time-calibrated tree are available in Dryad Digital Repository (#will be submitted as soon as the paper became accepted).

## Acknowledgments

We thank Andrea Cocucci, Alicia Sérsic, Birgit Harter Marques, Clemens Schlindwein, Reisla Oliveira, Eduardo Almeida, Gabriel Melo, Jerome Rozen, Juan Pablo Torreta, Mabel Lizaraso, Kelli Ramos, Leopoldo Alvarez, Rafael Ferrari, Rodrigo Gonçalves, and Vicent Lee for providing specimens for this study. We also thank Lilian Giugliano, Kelli Ramos and Anahí Espíndola for suggestions on previous versions of this manuscript. We thank P. Cedro for designing figures, M. Cavalcante, and V. Silva for helping with R programming and F. Duque for grammar revision. TMAR received scholarships from Fundação de Apoio à Pesquisa do Distrito Federal (FAPDF) and ACM received scholarships from CAPES and CNPq during the development of this research. AJCA thanks CAPES and FAPDF for grant support (process n. 193.000.893/2015).

